# Elasmobranchs’ metabarcoding requires a pragmatic approach to reach its promises

**DOI:** 10.1101/2022.08.25.505299

**Authors:** Marcelo Merten Cruz, Thomas Sauvage, Anthony Chariton, Thales Renato Ochotorena de Freitas

## Abstract

Human impacts have been eroding marine ecosystems in such a way that biodiversity patterns are changing. Therefore, policies and science-based solutions are indispensable for monitoring threats to the most impacted species. In such effort, the analysis of elasmobranchs’ environmental traces via eDNA metabarcoding represent a candidate tool for effective monitoring and conservation that is often advocated to be cost-effective and easily replicated. Here, we tested a realistic approach to monitor future changes through elasmobranchs’ metabarcoding with published primers, in which, elasmobranch diversity from the coastal waters of the Fernando de Noronha Archipelago (Brazil) was studied here. We detected a total of three elasmobranch species, namely *Hypanus berthalutzae, Ginglymostoma cirratum*, and *Prionace glauca* among numerous other fish species. Even though the technique proved to be a useful tool, some practical constraints were identified, and primarily caused by currently published environmental primers. In order to ensure the broad application of the method, we pointed out feasible adjustments to the problematic parameters based on our survey and other elasmobranch metabarcoding studies. The current drawbacks of the approach need to be considered by managers, conservation actors, and researchers, who are considering this methodology in order to avoid unrealistic promises for the cost incurred.

## Introduction

In the 21^st^ century, the ocean has been facing human-induced threats that are affecting the marine ecosystem in such a manner that extinction rates are dramatically increasing (Luypaert et al., 2020: Sage, 2020). Several elasmobranch (sharks, rays, skates, and sawfish) populations are declining and changing their distribution due to the wide variety of threats that they have been facing (Dulvy et al., 2014; 2017; 2021). For example, more than three-quarters of tropical and subtropical coastal species are threatened (Dulvy et al., 2021). Overfishing is the main threat for all threatened elasmobranch species evaluated by Dulvy et al. (2021). The ever-increasing consumption of fish in recent decades is also affecting the amount of elasmobranchs’ prey species (Navia et al., 2017). Elasmobranchs face additional issues such as fin trade, pollution, biomagnification, bycatch, and habitat loss (Pacoureau et al., 2021). Unfortunately, their naturally slow reproductive characteristics exacerbate these threats and hamper population recovery (Field et al., 2009).

The loss of elasmobranchs’ biodiversity may lead to irreversible environmental collapses because they (as high-level predators) help regulate and maintain the balance of marine ecosystems by structuring the dynamics of food webs (Navia et al., 2017). Considering it, there is a particular concern in establishing defined geographical areas for their protection (Stevens, 2010). To this extent, Marine Protected Areas (MPAs) are the main tool used in spatial management (Rigby et al., 2019). The establishment of MPAs is a common policy strategy to provide a refuge against anthropogenic-related threats and is considered vital to ocean health and resilience (Crear et al., 2020).

That is because well-managed MPAs can act as a recovery space, where species of particular importance, such as fishes, substantially increase in size, density, species richness, and biomass (Lester et al., 2009). When a long-term integral protection is implemented in a large area that captures a range of environments, the abundance, and diversity of fish populations can increase (Davies et al., 2021). To effectively reduce the disturbance to elasmobranch species within MPAs it is thus necessary to investigate their presence, distribution, and critical habitats within the protected range (Rigby et al., 2019). In such efforts, eDNA metabarcoding, an innovative molecular approach that allows identifying species presence from water samples to track the genetic material released by species in their environments, was previously developed to target elasmobranchs with specific primers (Bakker et al., 2017). Using this noninvasive technique, some authors have proven that more elasmobranch species can be identified than through traditional surveys (Boussarie et al., 2018). Using such approach, Bakker et al. (2017) were also able to demonstrate that elasmobranchs’ species diversity and eDNA read abundance directly reflects the known degree of marine anthropogenic impact.

The eDNA application gained a promising avenue when sequencing technologies improved the capacity to analyze fragmented genetic traces (Guardiola et al., 2015; Fraija-Fernández et al., 2020). With the aid of recent advancements, elasmobranchs’ biomonitoring studies rapidly advanced from the eDNA detection of target species (Simpfendorfer et al., 2016; Gargan et al., 2017; Weltz et al., 2017) to the differentiation between species through high-throughput sequencing - metabarcoding (Miya et al., 2015; Bakker et al., 2017; Boussarie et al., 2018; Collins et al., 2019). The metabarcoding approach has been proven to be an efficient tool for marine monitoring and ecosystem health assessment (Thomsen et al., 2012; Leray and Knowlton, 2015). And, recent studies provided a proof-of-concept that sharks, rays, and their relatives can be monitored using eDNA metabarcoding (Fraija-Fernández et al., 2020; Ip et al., 2021; Mariani et al., 2021; Marwayana et al., 2021; Monuki et al., 2021; Liu et al., 2022; Dunn et al., 2022).

In the present case study, we investigated the reliability of currently published elasmobranchs’ metabarcoding primers (Taberlet et al., 2018) to monitor the presence of this taxonomic group in a remote and resource-limited MPA known as Fernando de Noronha archipelago (FNA) in the equatorial Atlantic. FNA represents an important hotspot of biological diversity and a refuge for many species that have evolved on these islands through isolation (UNESCO, 2021). Choosing FNA as a model also bears the goal of evaluating the approach as a baseline for future policy implementation by the Brazilian Ministry of Environment to better protect and monitor the archipelago. Indeed, nearly 70% of the archipelago is designated as a “Marine National Park”, and the remaining areas as “Environmental Protected Area” (where some sustainable use of biodiversity is allowed) (Brasil, 1988).

Elasmobranchs are one of the main groups of FNA’s fish populations. Indeed, historical records indicate that it shelters at least 25 elasmobranch species (Soto, 2001; Garla and Garcia 2008). The use of FNA as a nursery habitat was reported for at least three shark species: *Carcharhinus perezi, Ginglymostoma cirratum*, and *Negaprion brevirostris* (Garla et al., 2005, 2009), which have been well-studied by acoustic telemetry and underwater observations (Garla et al., 2005; 2009; 2017a; 2017b). Nonetheless, habitat use and the diversity of migrating elasmobranchs species in the region are not fully understood, as well as the possibility of the site representing a nursery area for other species (Aguiar et al., 2009, Bucair et al., 2021).

Metabarcoding is often recommended as a cost-effective and easily replicated technique that could potentially be the answer to this and other monitoring gaps (Miya et al., 2015; Marwayana et al., 2021; Monuki et al., 2021). However, the elasmobranchs’ metabarcoding needs validation in realistic situations before being guaranteed as a routine monitoring tool. Here, the challenges to using this as an MPA monitoring tool were tested by using a case study of FNA. Our goal is to discuss if the elasmobranchs’ metabarcoding is mature enough to be considered as a routine tool for the monitoring of MPA’s and their management.

## 2. Materials and Methods

### 2.1. eDNA sampling

Collections were made in November 2020 under research permit No. 62360 from the Brazilian Ministry of Environment (SISBIO). The approach was a representation of localities across the north-western seascape of the Fernando de Noronha’s main island, as this was previously classified as an important area for sharks (Garla et al., 2009). Eight “sampling stations” were surveyed in triplicate resulting in 24 environmental samples. The sites were accessed by boat, in which 2 L of surface water was collected at each replicate by using sterile whirl-pack bags. A different pair of disposable gloves were used in each site to avoid eDNA cross-contamination. During the expedition, samples were kept in a thermal box up to 12h before filtration. This condition was purposely simulated because this is the realistic preservation condition that a remote MPA manager with a small boat could assume. In an improvised water filtration laboratory on land, each sample was individually filtered, through a Cellulose Nitrate filter (Millipore, 47 mm diameters and 0.45 μm pore size) using a vacuum pump and subsequently stored at room temperature in sterile microcentrifuge tubes filled with lysis buffer (TrisHCl 0.1 M, EDTA 0.1 M, NaCl 0.01 M and N-lauroyl sarcosine 1%, pH 7.5–8). The filtration apparatus was rinsed with commercial mineral water and sterilized with a 10% bleach solution and rinsed thoroughly between each sample. Two liters of commercial mineral water were used as the negative control and processed in the same way to the eDNA samples.

### 2.2. eDNA extraction, amplification, library preparation, and sequencing

All samples were transported to a quarantine facility within the LACE Lab (UFRGS, Porto Alegre/Brazil) where filters were stored at –20°C until further steps. Total eDNA was extracted using the DNeasy PowerWater Kit (Qiagen) according to the manufacturer’s protocol. DNA extracts were sent to EcoMol Consultoria where amplification, library preparation and sequencing steps were conducted. The performance of an elasmobranch-specific primers (targeting 170-185 base pairs of the 12S mitochondrial rRNA gene (“Elas02_F” 5’-GTTGGTHAATCTCGTGCCAGC-3’ and “Elas02_R” 5’-CATAGTAGGGTATCTAATCCTAGTTTG-3’, Taberlet et al., 2018) was first assessed against tissue-derived DNA extractions from sharks and rays commonly found in the Brazilian waters (samples from Merten Cruz et al. 2021). After confirming the successful amplification of shark/ray DNA extract, all collected eDNA samples were amplified with Elas02_F, and Elas02_R tailed with Illumina adapters (Nextera XT DNA Library Preparation Kit). PCR was carried out using PCR Biosystems Ultra polymerase mix (PCR Biosystems) in 25 μl reactions containing 250 nM of each primer and 2 μl extracted eDNA. Thermal conditions consisted of an initial denaturing 15 min at 95 °C, followed by 30 cycles at 95 °C for 30 s, 59 °C annealing for 30 s, 72 °C extension for 30 s, and a final extension of 5 min at 72 °C. Three PCR replicates per sample were done and subsequently pooled. A muscle tissue of *Rhizoprionodon terraenovae* (not reported in FNA) was also included as the PCR positive control. Also, negative controls were included for each PCR performed, to guarantee that no contamination occurred during the amplifications. Amplification was checked using 1.5% agarose gel (electrophoresis). PCR products were purified with magnetic beads (Agencourt AMPure XP® – Beckman Coulter) following the manufacturer’s instructions. After purification, a second PCR was performed with Nextera Index kit® adapters (Illumina). In this reaction, the adapters used as primers (P5 and P7) were compatible with the next-generation sequencing Illumina system. A unique combination of the index was used (specific sequences associated with the Illumina adapter) to identify each sample further. Later, the second amplification products were evaluated on 1.5 % agarose gel. Finally, the second PCR products were again purified with magnetic beads (Agencourt AMPure XP® – Beckman Coulter), quantified using NanoDrop® 2000 (Thermo Scientific), and adjusted to 20ng/μl. Then, samples were equimolarly pooled and purified by electrophoresis to remove spurious fragments much smaller than the desired fragment size. The purified pool was quantified by real-time PCR with the Kapa Biosystems Quantification Kit (Roche) according to the manufacturer’s protocol. The final libraries were sequenced on an Illumina iSeq® platform using the Illumina kit iSeq 100 i1 V2 300cycles (2×150bp paired-ends) at a final concentration of 150 pM. The extraction, pre-PCR, and post-PCR procedures were all carried out in physically separated rooms. PCR steps were conducted under the hood using exclusive micropipettes previously disinfected under 15 minutes of UV light. Materials and surfaces were cleaned using a 10% bleach solution.

### 2.3. Bioinformatics and Data analysis

After sequencing, the quality of raw reads was checked using the FastQC software using base quality scores >30. Sequences were then processed in a custom pipeline in Rstudio Software (v4.0.2; R Studio Team, 2021). The cutadapt package was used to remove the forward and reverse primers from the reads, allowing a maximum of two errors on primers (Martin, 2011); Then, the sequences were processed in a custom pipeline using the DADA2 (version 1.6; Callahan et al., 2016) in which, paired-end reads were merged and sequences filtered to keep those longer than 225 bp and eliminate those shorter than 130 bp. Chimeras and reads containing indeterminate bases (Ns) were also removed. The remaining reads (i.e. Amplicon Sequence Variants, ASVs) were kept for downstream taxonomy assignment. ASVs occurring with less than 50 reads were removed. The resulting sequences were subjected to taxonomic identification using the assignTaxonomy function provided by DADA2. The first pass taxonomic assignment was queried against the customized reference library (Table Supll.1) with an identity score greater than or equal to 98%. To further investigate the marine vertebrate fauna in FNA, a second taxonomic assignment using BLASTn against the NCBI nucleotide database (accessed in January 2022) using the percent alignment identity as search parameters. Only sequences with an identity higher than 95% with a sequence from the reference database and classified as marine vertebrates were considered for further analyses. A sample-based species accumulation curve was also calculated to estimate sampling effort using the R package vegan 2.5-7(Oksanen 2022). To do so, the sum of the marine vertebrates’ sequence read counts corresponding to each sample was plotted as a function of the number of samples (Fig. 2).

## Results

The eDNA amplicons yielded a total of 1,311,609 reads, including the positive controls as well as amplification and sampling negative controls that were removed after the applied pipeline. The positive control sequences corresponding to *Rhizoprionodon terraenovae* was successfully recovered. After bioinformatic filtering, 85,313 merged, paired-end, non-chimeric, and assembled reads were obtained. Of these, 23,105 belonged to marine vertebrate species. This number represents only 1.51% of the total reads, as most of the reads corresponded to non-specific amplicons. The Elas02 primer pair appeared to amplify a longer section of the bacterial 16S rDNA, whose length did not allow pairing with their reverse complement. These 16S contaminants, being off-target (i.e., not elasmobranchs or metazoa), were excluded from subsequent analysis.

The targeted survey for elasmobranchs resulted in three identified species with 100% of identity matched in the reference database. *Hypanus berthalutzae* was detected in six out of the eight sampling stations, resulting in a total of 3,470 merged reads among all samples (15% of the filtered reads), while *Ginglymostoma cirratum* could be surveyed in three sites; 626 merged reads among all samples were taxonomically assigned to this species (2.71% of the filtered reads). Besides, *Prionace glauca* was present in two sites, and 662 merged reads out of the total corresponded to this shark (2.86% of the filtered reads).

Aside from the elasmobranchs’ identifications, the GenBank BLASTn analysis identified 37 other marine vertebrate species. Most sequences matched their target with a very high identity level (>95%). However, four species (*Hygophum benoiti, Hyporhamphus yuri, Pempheris molucca, Chriolepis dialepta*) were excluded from the subsequent analysis as their GenBank database BLAST matches were below 95%. One cetacean species was observed: the spinner dolphin (*Stenella longirostris*), which represented 5.85% of the retained reads. All the other marine vertebrate species identifications belonged to the bony fishes (73.55% of the retained reads). The final number of bony fishes’ assignments was then 32 species in 17 families (Table 1). The “redtail parrotfish” (*Sparisoma chrysopterum*) was the most represented species in this survey, comprising 53.20% of the marine vertebrate reads.

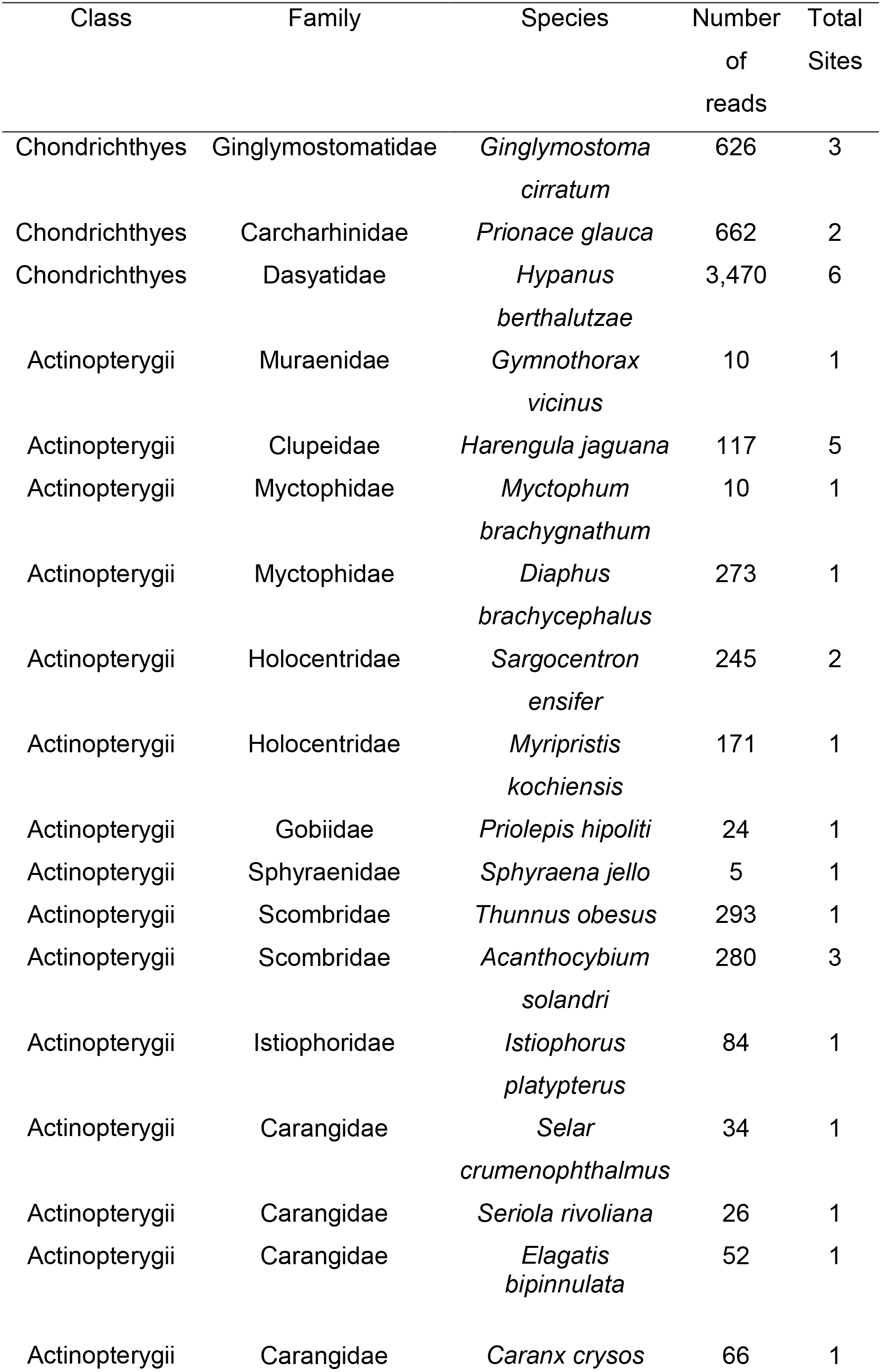
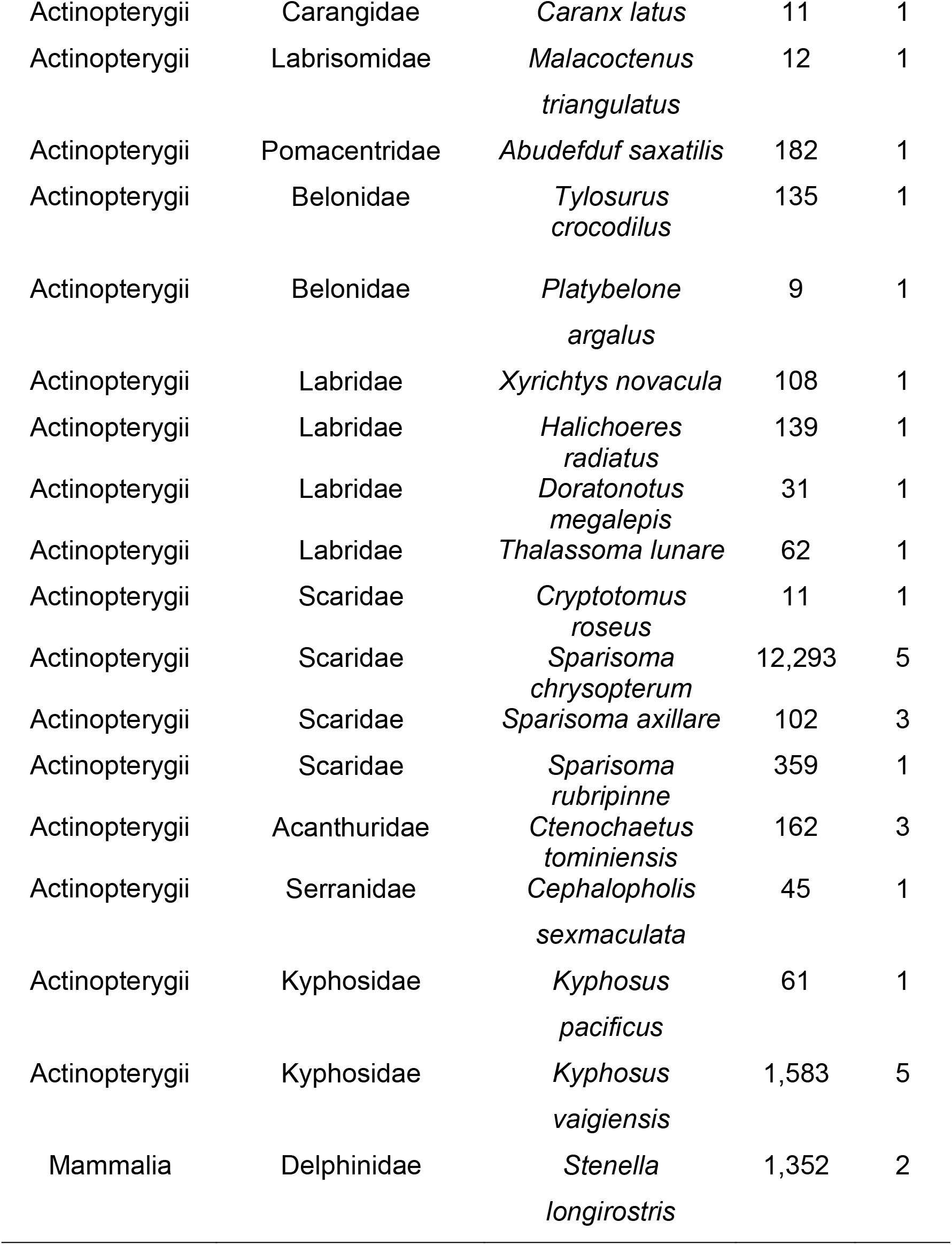

The average number of marine vertebrates’ sequences per sample was 963, which was insufficient to characterize biodiversity patterns across the sampled sites. The most diverse site was sampling station number 1, corresponding to the region near the port of the main island (Figure 1). The species accumulation curve plateaued after 10 and 15 eDNA samples, indicating that augmenting the sampling effort would not necessarily increase the number of species detected with the primer used (see further above regarding lack of specificity issue).

**Figure 1:**
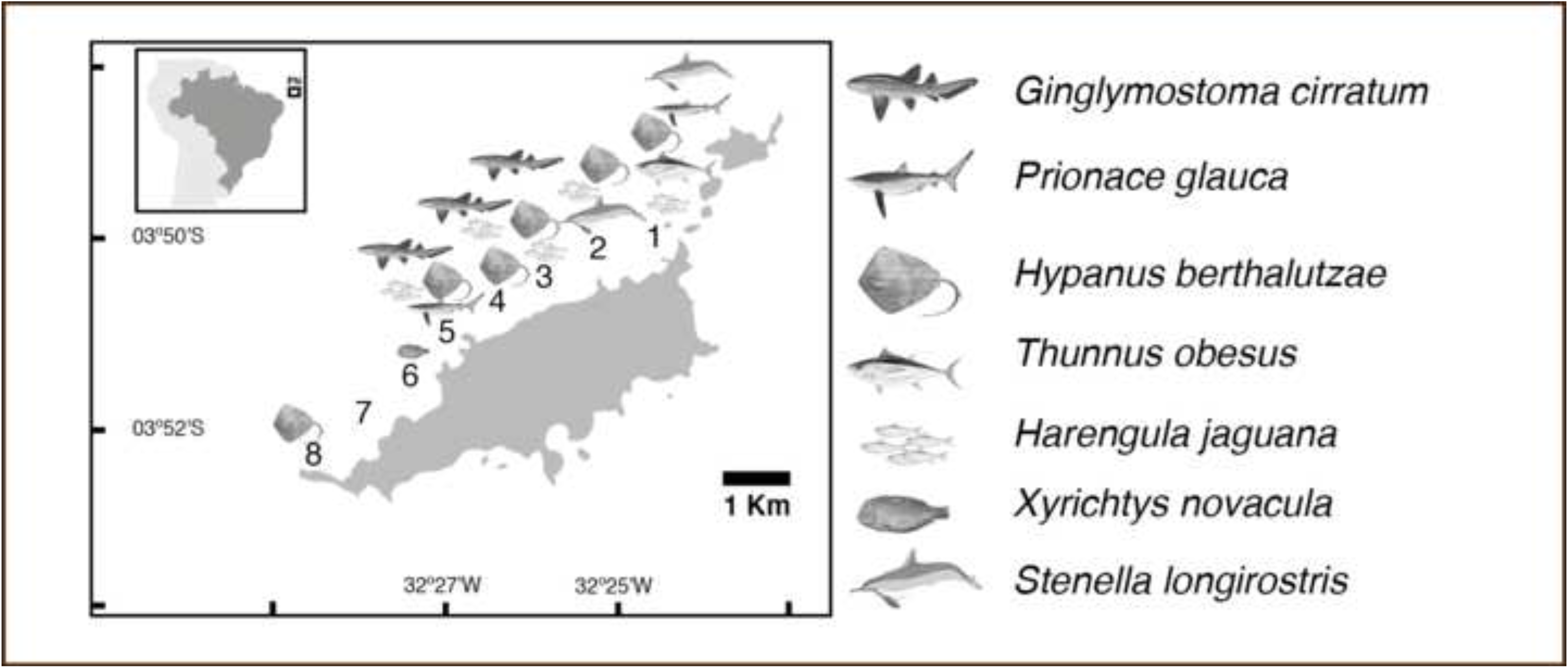
Distribution of the major findings (*Ginglyomostoma cirratum, Prionace glauca, Hypanus berthalutzae, Thunnus obesus, Harengula jaguana, Xyrichtys novacula*, and *Stenella longirostris*) across the northwestern coast of Fernando de Noronha Archipelago.

**Figure 2:**
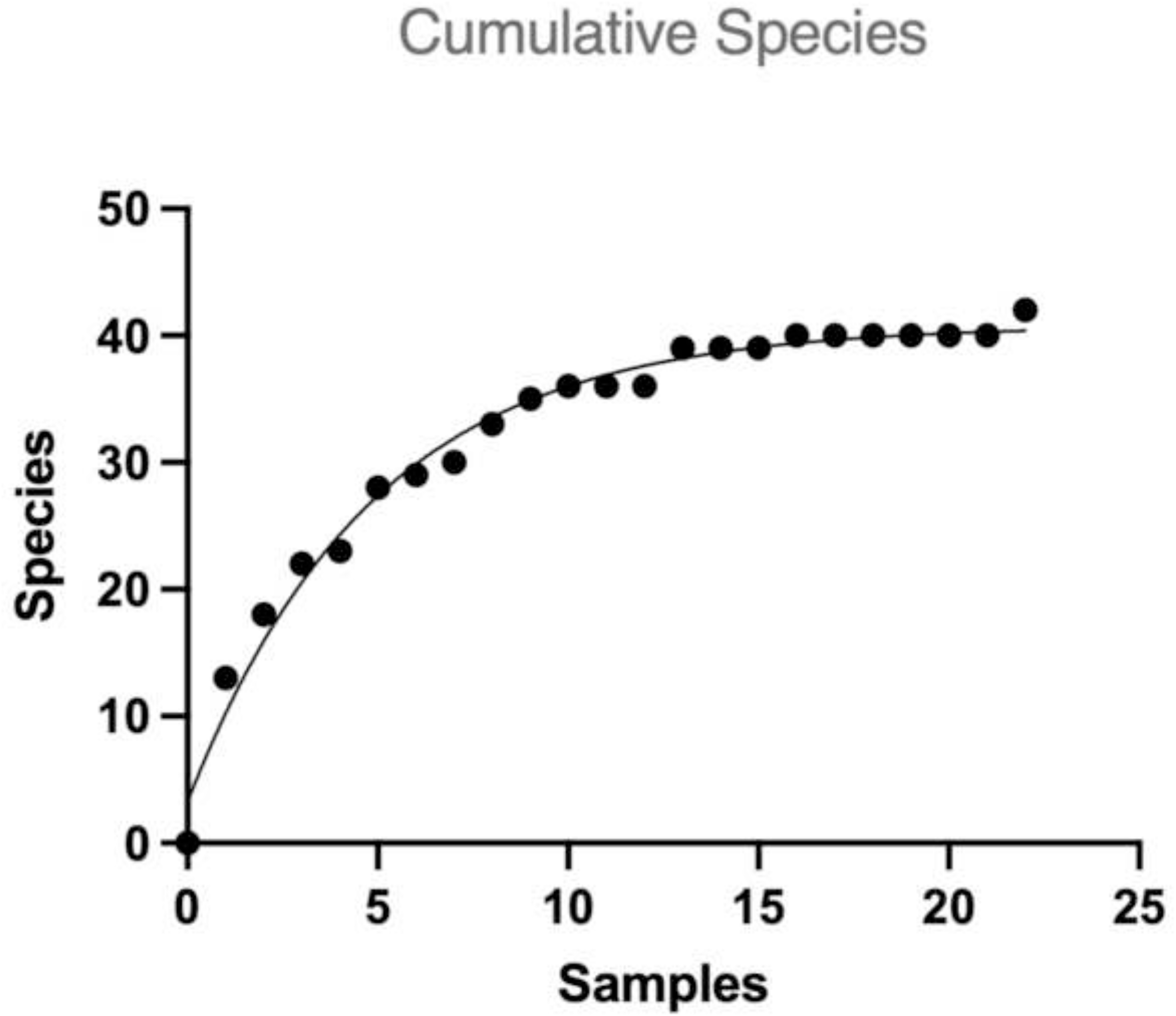
Accumulation curve of identified species in water samples from the northwestern coast of Fernando de Noronha Archipelago.

## Discussion

Elasmobranchs are on the brink of extinction, and due to the lack of data, they are facing a troubled future. In this context, fine-tuning biomonitoring tools may represent a valuable approach to better understanding elasmobranch populations and their threats. Since oceanic areas are vast, study models’ sites and representative species are needed to predict changes. The best model choice will potentially be informative, robust, replicable, and cost-effective. The application of metabarcoding for elasmobranchs’ monitoring is maturing fast as an adequate tool for those demands. However, our results with existing environmental primers demonstrate that engaging in a pragmatic approach is still necessary before this tool can be endorsed as a routine monitoring tool.

### Metabarcoding at Fernando de Noronha

This survey contributes to the increase of knowledge of the recently identified *Hypanus berthalutzae* (Petean et al., 2020). This stingray species is endemic to Brazilian waters, yet little is known about its ecological aspects; even so, their presence in habitats closer to the coast and in shallow environments is confirmed here (Branco-Nunes et al., 2021). Given that metabarcoding-based biodiversity assessments have proven to be a powerful method in occupancy modeling (McClenaghan et al., 2020); the heterogeneous preferences of *Hypanus berthalutzae* in shallow sandy and reef areas in the Fernando de Noronha archipelago could be further investigated using this application (Branco-Nunes et al., 2021).

Additionally, the applied methodology detected the “blue shark” (*Prionace glauca*), an elasmobranch previously reported in the archipelago; however, their habitat use in the region has not yet been investigated (Soto, 2001). The blue shark is pelagic; its distribution is the widest among all large sharks, and it is common throughout the coast of continental Brazil (Hazin and Lessa, 2005). Due to their highly migratory nature, their presence in Fernando de Noronha’s waters is probably occasional (Hazin et al., 1994). In this regard, eDNA analysis has been gaining traction in the surveillance of migratory sharks, and such use could improve temporal and spatial data for management purposes (Postaire et al., 2020; Schweiss et al., 2020).

Also, the detection of *Ginglymostoma cirratum* in its described nursery ground (Garla et al., 2009) is constructive to the application of this tool in the identification of critical habitats, as the *Ginglymostoma cirratum*’s eDNA could be uncovered due to their high-site fidelity, which may increase the eDNA signal and facilitate molecular detection (Garla et al., 2019; West et al., 2020).

Finally, Mariani et al. (2021) pointed out that metabarcoding surveys can offer the “bonus” of non-target vertebrate sequences. Additional biodiversity could be catalogued in this manner. For example, the spinner dolphin (*Stenella longirostris*) is especially relevant because this species is one of the symbols of Fernando de Noronha and is important for tourism, public engagement, and, more broadly, biodiversity conservation (Tischer et al., 2020). The survey allowed for the identification of teleost biodiversity; both reef fish and pelagic (open water) species were recovered. The metabarcoding approach confirmed the island as a biodiversity hotspot for jacks (*Seriola rivoliana; Elagatis bipinnulata; Caranx crysos; Caranx latus*) and parrotfishes (*Cryptotomus roseus Sparisoma chrysopterum Sparisoma axillare Sparisoma rubripinne*) (Mazzei et al., 2017; Schmid et al., 2020). Beyond that, the method captured molecular signatures from eels (*Gymnothorax vicinus*); sardines (*Harengula jaguana*); tunas (*Thunnus obesus*); wahoos (*Acanthocybium solandri*); sailfishes (*Istiophorus platypterus*); needlefishes (*Tylosurus crocodilus; Platybelone argalus*); and wrasses (*Xyrichtys novacula; Halichoeres radiatus; Doratonotus megalepis*).

However, our results suggest that the co-amplification of non-targeted species might represent a disadvantage. For instance, the incidental amplification of prokaryotic DNA by the Elas02 primer represents an important waste of sequencing depth/effort (Collins et al., 2019). As also reported by Dunn et al. (2022), the adjustment of some parameters may potentially increase the coverage of elasmobranch detection

### Challenges, lessons learned, and feasible solutions

Elasmobranchs’ metabarcoding has been reaching great outcomes reporting the successful detection of studied species, in particular when compared to conventional biomonitoring methods (see Boussarie et al., 2018; Ip et al., 2021; Liu et al., 2022); nonetheless, the species undetected by eDNA remain poorly discussed. As metabarcoding is still a growing field, it is impossible to know if the absence of expected species is true or if it results from methodological limitations (Liu et al., 2022; Dunn et al., 2022). Here, we discuss the potential increase in the detection rate of elasmobranch species (Box 1).

The main constraint in capturing elasmobranchs’ diversity in the Fernando de Noronha case study was the incidental amplification of non-target DNA (waste of sequencing effort). Once bacterial DNA became overabundant in our samples, the low specificity of the Elas02 primer cross-amplified prokaryotic sequences rather than elasmobranchs’ (interference phenomena explained by Wilcox et al., 2013). In detail, the mitochondrial 12S rRNA gene shows several regions of significant homology to bacterial 16S rRNA sequences (Eperon et al., 1980). Previous metabarcoding studies faced the issue of cross-reactivity of a metazoan 12S rRNA primer with bacterial 16S rRNA sequences (Machida et al., 2012; Lim and Thompson, 2021). The opposite also occurs (Huggins et al., 2020). Most recently, Dunn et al. (2022) further corroborate that elasmobranch 12S primers should be improved or redesigned as they are inefficient for the amplification of elasmobranch eDNA.

Nevertheless, Collins et al. (2019) tested a variety of elasmobranch primers and concluded that 12S performed better than COI in terms of specificity, discriminatory power, and reproducibility for this specific taxonomic group detection. Non-target amplification also occurred when Bakker et al. (2017) applied a COI marker to infer shark presence in tropical habitats. According to Siddall et al. (2009), marine prokaryote sequences are presumed to be amplified by metazoan COI primers due to nucleotide convergence. Dunn et al. (2022) concluded that further studies into the targeted elasmobranchs’ metabarcoding primers would be required to maximize shark and ray monitoring outcomes. Thus, one option is to use both markers side by side in a multi-marker approach to maximize elasmobranch biodiversity recovery (see Ip et al., 2021). Another alternative can be to design a specific primer to amplify the elasmobranchs of a given region using a regional-specific reference library (Taberlet et al., 2018). The Supplementary Database can be used as a baseline for future metabarcoding studies in FNA. Developing an elasmobranch-specific primer may be difficult. Utilizing universal fish primers may be a better approach to detecting sharks within the fish population (see Fraija-Fernández et al., 2020).

Although the Elas02 marker has not been shown to be a good choice and certainly the alternatives mentioned above must be considered, other studies have reached expressive results using the same primer (Mariani et al. 2021; Liu et al., 2022). Then other methodological changes should be investigated. For example, it is known that eDNA degrades relatively quickly in marine systems (Collins et al., 2018). Particularly, high temperatures and ultraviolet (UV) radiation can decrease the eDNA signal (Barnes et al., 2014; Andruszkiewicz et al., 2017; McCartin et al., 2022). It is possible that this occurred due to the implementation of a realistic sampling strategy – similar to what MPA’s managers in tropical areas would use (i.e., lack of refrigeration, logistical constraints, and prolonged periods of time before the filtering procedure). To overcome this challenge when refrigerators are infeasible, it seems a plausible recommendation to substitute Whirl-Pak bags in a rigid and hermetic sampler (for example, Nalgene Sample Bottle, used by West et al., 2020; Liu et al., 2022) and protect eDNA samples cooled and stored on ice until filtering, similarly as described by Simpfendorfer et al., (2016) when rare species in a logistic-limited environment are detected.

One of the main recommendations to better estimate fish diversity is to increase the number of samples (Juhel et al. 2020; Marwayana et al., 2021). By analyzing our data, the accumulation curve rarefied with fewer than 15 samples, so it is possible to assume that a greater sampling intensity would not reflect more elasmobranchs being discovered on the NW coast of FNA using our strategy. Instead, the low number of targeted reads indicates which parameters should be optimized, focusing on the integration of metabarcoding into the MPA framework. Based on the research of Boussarie et al. (2018), which successfully assessed shark biodiversity in an isolated archipelago with 22 samples, a restricted number of samples is a reliable parameter to maintain when working with oceanic islands.

In essence, capturing DNA fragments from elasmobranchs in seawater can be particularly challenging, as they usually occupy the upper-level trophic position, so naturally, their density is reduced compared to species occupying lower levels (Postaire et al., 2020). Moreover, elasmobranchs’ populations are in decline and consequently, these species are a difficult target as the density will potentially reflect the sensitivity of eDNA detection (Furlan et al., 2016; Adams et al., 2019). The authors recognized that 2 L of the environmental sample in an open-water habitat is insufficient to capture elusive DNA. In turn, Bakker et al. (2017); Boussarie et al. (2018); and Mariani et al. (2021) achieved their goals by analyzing 4 L of seawater. In reality, maximizing water volumes as much as logistically possible will potentially increase fish eDNA detection (Bessey et al., 2020).

Additional sampling strategies are required to find elasmobranchs’ eDNA to circumvent their habitat characteristics. For example, elasmobranchs are often highly migratory, so their presence in a specific environment is ephemeral as their eDNA is released (Schweiss et al., 2020). Likewise, FNA’s elasmobranchs follow seasonal movements (Garla et al., 2005, 2009; 2019); therefore, a monthly sampling approach should be established to track transient occurrences and changes in elasmobranchs’ biodiversity throughout the seasons (Postaire et al., 2020). Furthermore, some elasmobranchs are highly associated with the seafloor and the limiting condition that water column stratification restricts vertical eDNA transport, is a huge pitfall for the surface sampling methodology (Jeunen et al., 2020). Considering that, other strategies must be examined. For instance, Ip et al. (2021) effectively enhanced the precision of elasmobranch diversity estimates by analyzing sediment environments. Another procedure to be considered is the replicating of the samples from varying standardized depths, as Marwayana et al. (2021) suggested. Also, a horizontal sampling intensification varying across habitats will be required to completely study heterogeneous ecosystems (Jeunen et al., 2019; West et al., 2020; Marwayana et al., 2021).

Lastly, our results highlighted the importance of a regional-specific reference library to bio-surveillance purposes as a general recommendation. For example, the non-target marine vertebrates’ taxonomic assessment using the NCBI database revealed some “closest match” species not associated with FNA’s surroundings (*Myctophum brachygnathum, Sargocentron ensifer, Myripristis kochiensis, Sphyraena jello, Thalassoma lunare, Ctenochaetus tominiensis, Cephalopholis sexmaculata*, and *Kyphosus pacificus*). The detected traces are probably from FNA’s species that are phylogenetically closer (i.e., low differentiation between 12S sequences) to those misidentified such as *Myripristis jacobus, Sphyraena barracuda, Thalassoma noronhanum, Cephalopholis cruentata* or *Cephalopholis fulva*, and *Kyphosus vaigiensis*, respectively (Soto, 2001).

### Considerations for the broad application of metabarcoding methods to monitor elasmobranchs

The metabarcoding monitoring field continues to mature, and it is natural that improvements will appear as much as the methodology is applied. What should be considered with responsibility is that almost all amendments raised above could potentially hamper the two main eDNA promises: to be cost-effective and repeatable (Evans et al., 2017; Dickie et al., 2018).

Marine eDNA technologies are frequently pointed out as a cost-effective tool, mostly because it is more economical than underwater visual census (UVCs) and baited remote underwater video stations (BRUVS) operations (Boussarie et al., 2018; Stat et al., 2019; Ip et al., 2021). Future investigators should be aware that a metabarcoding study is not uncostly. Moreover, metabarcoding research is an expensive investment for middle and low-income economies, for example, all other studies listed in Box 2 were led by countries classified by the World Bank as a “high-income economy” (The World Bank, 2022). Methodological challenges are imposed on scientists who are not from wealthy countries, for example, the exchange rate from dollars or euros to home currencies that is often unfavorable (Valenzuela-Toro and Viglino, 2021). Then, as the extraction, amplification, and sequencing materials are usually dollar-priced by international companies, an intensification of sampling in metabarcoding surveys is not practical in economic turmoil realities.

Without diversifying the elasmobranchs’ metabarcoding narrative, scientists from developing and least developed countries are imperiled to be passive spectators rather than active contributors to this revolutionary method (Ahmadia et al., 2021). Statements such as, “Additional samples could be taken from varying depths without adding significant field costs” (Marwayana et al., 2021); and “given the ease and cost-effective nature of eDNA sampling, such sampling efforts are not cost prohibitive” (Monuki et al., 2021); are susceptible to offering and unrealistic expectation to those researchers whose science-investment choices are limited.

The high risk of contamination is one of the main drawbacks of using water samples to capture molecular signatures of aquatic species, and careful considerations were already raised (Sepulveda et al., 2020). Indeed, metabarcoding’s strict protocols are demonstrated to be mandatory, as seen from the overabundance of “non-fish” DNA we obtained when testing relaxed conditions. Ideally deliberate considerations are demanded to avoid contamination in marine sampling, so it would be necessary to have equipped boats, a trained crew, and a dedicated molecular laboratory on land (Goldberg et al., 2016; Gilbey et al., 2021). Those conditions are often minimized in proponent studies or presented without weighing the real difficulty involved in incorporating actions such as collecting water with a crane system from a boat (proposed by Marini et al., 2021); or using the continuous circuit intake of a ship (Fraija-Fernández et al., 2020); or keeping samples at − 80 °C in less than 2 hours between the collection and the storage (Ip et al., 2021).

Furthermore, logistical aspects should be considered when a methodology is recommended (Dickie et al., 2018). For example, sampling strategies that maximize the detection of elusive species are certainly important, but the feasibility of their implementation by MPAs is not fully discussed; such as the real challenge of assessing 40 m of depth in a survey (proposed by Marini et al., 2021; Dunn et al., 2022); or to collect multiple samples of 100 L (Valentini et al., 2016); or to collect more than 250 1L samples in an expedition (West et al., 2020).

Those are some impediments to incorporating metabarcoding in remote and limited-resources MPAs. Such restrictions will naturally cause a social gap between who is able to use such technology and consequently a detachment of fishers, managers, community members, and local people in elasmobranch conservation (Rigby et al., 2020). In order to approximate managers and monitors of eDNA tools, the non-technical summary proposed by Gilbey et al. (2021) is the appropriate guidance. Paradoxically, this document is not freely available to non-specialists; this indicates that inaccessibility is still a challenge, and further discussion about open-science practices in conservation biology is necessary (Roche et al., 2022).

## Conclusion

Elasmobranchs are on the frontline of anthropogenic pressures, and future changes in their biodiversity patterns can act as a model of human impact management. Metabarcoding has been emerging as a solution tool for sensible monitoring of those changes. Even though the application of the technique brings significant outcomes to elasmobranch research, the optimal balance between gold-standard protocols and the feasibility of its implementation in some MPAs is still a research topic. For example, we could detect elasmobranchs through metabarcoding in a study case MPA, but not without limitations. An intensification of sampling and sequencing efforts allied with further development into the primers used could overcome these concerns. Nevertheless, the limitations in using the methodology in countries facing economic constraints must be weighed. Once that issue has been overcome, the method can be used in complementation with the more traditional techniques, which are preferable to “silver bullet” conservation actions, and researchers should put greater attention to reporting broadly applied reproducible field sampling methods.

## Acknowledgements

Concerning the samples, the authors would like to express their gratitude to the ICMBio Fernando de Noronha for their help and their support. This work was supported by the Federal University of Rio Grande do Sul (UFRGS), Macquarie University, the Brazilian Agency of the Coordination for Improvement of Higher Education Personnel (CAPES); the National Council for Scientific and Technological Development (CNPq), and the Research Support Foundation of the State of Rio Grande do Sul (FAPERGS).

## Authors Contributions

MMC, AC and TROF conceived and the study, MMC conducted the sampling, MMC and TS conducted all experiments, MMC and TS analyzed the data, MMC wrote the manuscript, AC, TS and TROF reviewed the manuscript, all authors read and approved the final version.

---

**Box 1:** Feasible adjustments to the elasmobranch metabarcoding’ constraints based on Fernando de Noronha’s survey and other elasmobranch metabarcoding studies

| Constraint | Feasible solution |
| --- | --- |
| Number of samples | It is possible to maintain a restricted number of samples; Good results with 22 samples (Boussarie 2018) |
| Avoid eDNA decay | To use Nalgene Sample Bottle (West et al., 2020; Liu et al., 2022), protect on ice until filtering (Simpfendorfer et al., 2016) and filter the samples as soon as possible (Barnes et al., 2014) |
| Incidental amplification of non-target DNA | Further research into the primers used (Dunn et al., 2022) |
| Capturing elusive species | To collect water as much as logistically possible (Bessey et al., 2020); but good results with 4 L (Bakker et al., 2017; Boussarie et al., 2018; and Mariani et al., 2021) |
| Capturing migratory species | To sample seasonally (Postaire et al., 2020) |
| Capturing benthic batoids species | To collect sediment environments (Ip et al. 2021) and/or to sample replications from varying standardized depths (Marwayana et al. 2021) |
| Studying heterogeneous ecosystems | To sample horizontally, varying across habitats (Jeunen et al., 2019; West et al., 2020; Marwayana et al., 2021) |
| General recommendation | To use a regional-specific reference library |

**Box 2:**
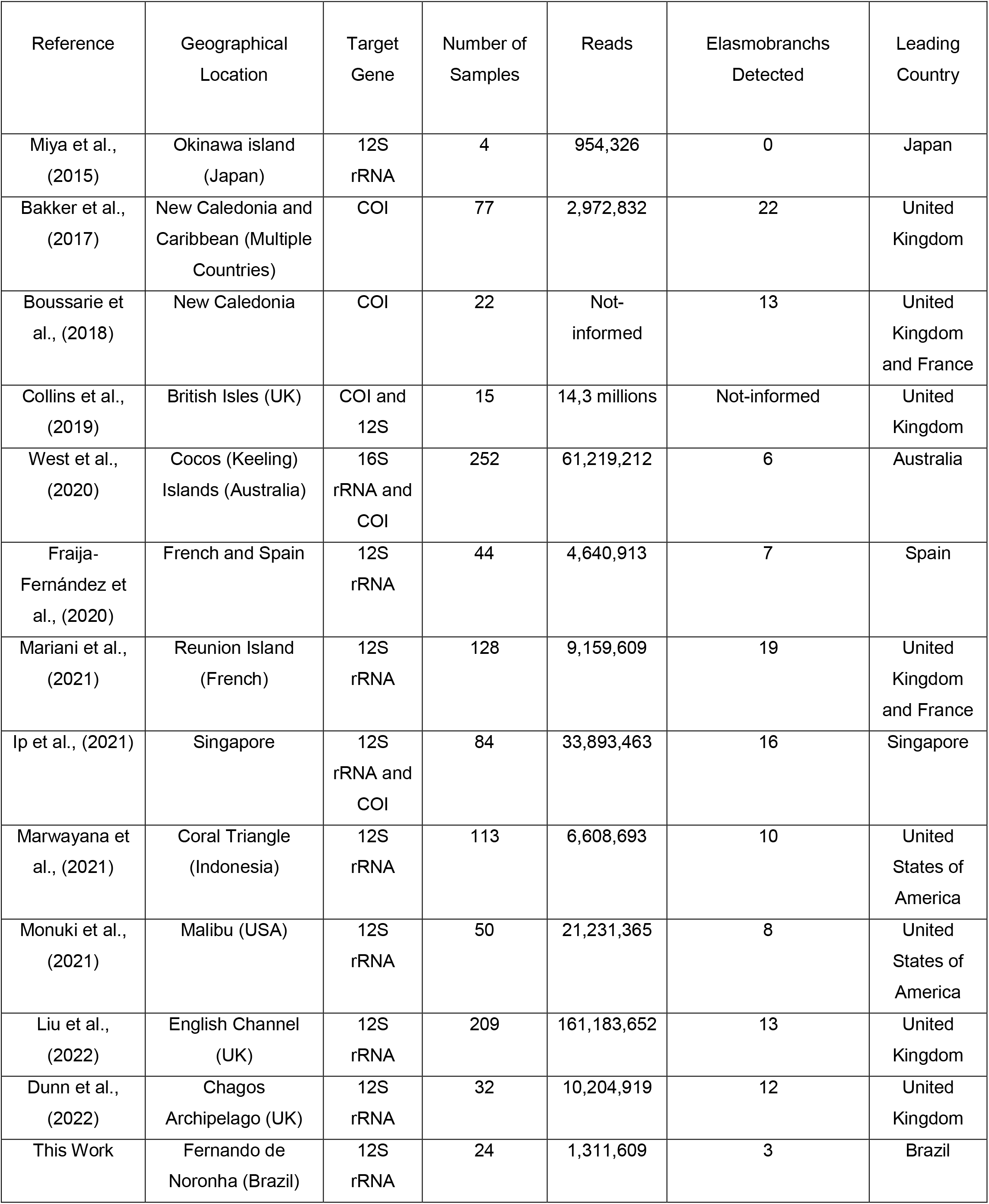
Compilation of Elasmobranchs’ Metabarcoding studies.

**Supplementary Table 1:**

Reads identification prior to removing unmerged reads to show the prokaryotic contamination.

**Supplementary Table 2:**

12S reference database from FNA’s Elasmobranchs

